# Persistent trajectory-modulated hippocampal neurons support memory-guided navigation

**DOI:** 10.1101/676593

**Authors:** Nathaniel R. Kinsky, William Mau, David W. Sullivan, Samuel J. Levy, Evan A. Ruesch, Michael E. Hasselmo

**Affiliations:** Center for Systems Neuroscience, Boston University, 610 Commonwealth Ave., Boston, MA 02215, USA; Graduate Program for Neuroscience

## Abstract

Trajectory-dependent splitter neurons in the hippocampus encode information about a rodent’s prior trajectory during performance of a continuous alternation task. As such, they provide valuable information for supporting memory-guided behavior. Here, we employed single-photon calcium imaging in freely moving mice to investigate the emergence and fate of trajectory-dependent activity through learning and mastery of a continuous spatial alternation task. We found that the quality of trajectory-dependent information in hippocampal neurons correlated with task performance. We thus hypothesized that, due to their utility, splitter neurons would exhibit heightened stability. We found that splitter neurons were more likely to remain active and retained more consistent spatial information across multiple days than did place cells. Furthermore, we found that both splitter neurons and place cells emerged rapidly and maintained stable trajectory-dependent/spatial activity thereafter. Our results suggest that neurons with useful functional coding properties exhibit heightened stability to support memory guided behavior.

## INTRODUCTION

Place cells in the hippocampus encode the current position of many different animals and humans (Ekstrom et al., 2003; Geva-Sagiv, Romani, Las, & Ulanovsky, 2016; Miller et al., 2013; Muller & Kubie, 1987; Muller, Kubie, & Ranck, 1987; Niediek & Bain, 2014; O’Keefe, 1976; O’Keefe & Dostrovsky, 1971) supporting the known role of the hippocampus in spatial memory and navigation across species (Morris, Garrud, Rawlins, & O’Keefe, 1982; Vorhees & Williams, 2014). However, the hippocampus is also widely known for its role in supporting the encoding, retrieval, and consolidation of non-spatial long-term memories (Corkin, 1984; Eichenbaum, 2004; Milner, Corkin, & Teuber, 1968), suggesting that it must represent variables beyond an animal’s current location. Indeed, recent studies have demonstrated that the hippocampus encodes the dimensions of a given task, from odors (Muzzio et al., 2009; Wood, Dudchenko, & Eichenbaum, 1999) to time (Howard et al., 2014; Kraus, Robinson II, White, Eichenbaum, & Hasselmo, 2013; MacDonald, Lepage, Eden, & Eichenbaum, 2011; Manns, Howard, & Eichenbaum, 2007; Pastalkova, Itskov, Amarasingham, & Buzsáki, 2008; Robinson et al., 2017; Salz et al., 2016) to tones (Aronov, Nevers, & Tank, 2017). One early demonstration that the hippocampus encodes dimensions beyond an animal’s current location was the discovery of trajectory-dependent neurons or splitter neurons (Frank, Brown, & Wilson, 2000; Wood, Dudchenko, Robitsek, & Eichenbaum, 2000), cells whose firing rate within a particular position was modulated based on the animal’s past or future trajectory in a spatial alternation task. The generation of this neural correlate suggests a potential mechanism allowing the hippocampal code to support both memory and decision based planning.

Several studies have demonstrated place cell firing fields move, or remap, their locations in response to new learning during a spatial learning task (Dupret, O’Neill, Pleydell-Bouverie, & Csicsvari, 2010; McKenzie, Robinson, Herrera, Churchill, & Eichenbaum, 2013). These studies highlight that the flexible adjustment of place field locations is important for learning new information. Conversely, the ability of hippocampal neurons to maintain the same firing location in the absence of learning might support long-term memory retrieval. In support of this idea, a recent study illustrated that neurons with place fields located near a hidden goal were more stable over time than cells with fields in other locations (Zaremba et al., 2017). Two other experiments found that increasing rodents’ attention to a task selectively heightened stability in neurons that encoded task-relevant features (Kentros, Agnihotri, Streater, Hawkins, & Kandel, 2004; Muzzio et al., 2009). These studies, along with the finding that place cells with fields in close proximity to a goal location exhibit heightened activity in post-learning sleep (Dupret et al., 2010), suggest that the utility of a neuron’s information to task performance influences its long-term stability.

Thus, since splitter neurons provide immediately relevant information for performing a spatial alternation task, we hypothesized that these neurons are important for successful task performance. Furthermore, we hypothesized that due to their utility, splitters may exhibit different long-term dynamics when compared to place cells. Specifically, we addressed three lines of inquiry. First, does the level of trajectory-dependent information within the hippocampus correlate with behavioral performance? Second, given the steady evolution of hippocampal activity patterns across days (Cai et al., 2016; Mau et al., 2018; Rubin, Geva, Sheintuch, & Ziv, 2015; Ziv et al., 2013), do splitter neurons remain part of the active population longer than other cells, thus providing a longer lasting memory or planning signal to guide behavior? Third, once a neuron establishes trajectory-dependent activity, is it less prone to remapping than other neurons? These questions are particularly relevant since trajectory-dependent activity has been observed in other tasks (Ferbinteanu & Shapiro, 2003; Smith & Mizumori, 2006b) and could be employed more generally by the hippocampus to guide the appropriate behavior based on environmental cues (Smith & Mizumori, 2006a).

To track neurons across long timescales, we paired a continuous spatial alternation task with *in vivo* miniscope recordings of GCaMP6f activity in dorsal CA1 of freely-moving mice. This technology allowed us to not only track the long-term activity of neurons, but also to adequately characterize the heterogeneity of trajectory-dependent activity in the hippocampus, since we can simultaneously record from a large number of neurons in each session. We first found that trajectory-dependent coding correlates with task performance, suggesting it is important for supporting memory-guided behavior. Second, we established that a neuron’s functional coding properties, also referred to as functional phenotype below, are important for predicting its long-term activity: splitter neurons were more likely to be persistently active in the days following their onset than were return arm place cells indicating that neurons which provide more adaptive information might provide longer lasting input to downstream structures. Third, we found that trajectory-dependent neurons display more consistent long-term information about an animal’s location than pure place cells. Fourth, we found that the population as a whole displayed a rapid onset of trajectory-dependent activity followed by stable coding of trajectory thereafter. Last, we discovered that recruitment of context-dependent splitter cells peaked several days into training, whereas place cell recruitment peaked on the first day. These results combined suggest that neurons which develop the most behaviorally important coding properties are preferentially stabilized in both their short and long-term dynamics, which enables them to more consistently and effectively support memory-guided behavior. Our research paves the way for future studies investigating how heterogeneity in the neural code might support acquisition and retention of more complex behavioral tasks.

## RESULTS

### Behavior and Imaging

Food deprived mice (n=4) with neurons expressing GCaMP6f in region CA1 of the dorsal hippocampus were trained to perform a continuous spatial alternation task on a figure-8 maze (Figure 1A) while we simultaneously recorded calcium activity using a miniaturized microscope. Mice exhibited a range of learning rates, taking from 5 to 21 sessions to acquire the task, which was defined as the third consecutive session of performance at or above our criteria of 70% (Figure 1B). Mice performed continuous alternation at or greater than criteria on average throughout the course of the experiment (Figure 1C). We utilized custom-written software (Kinsky, Sullivan, Mau, Hasselmo, & Eichenbaum, 2018; Mau et al., 2018) to extract neuron ROIs (Figure 1D), construct their corresponding calcium traces, and identify each ROI’s putative spiking activity (Figure 1E). Using this technique, we recorded from large numbers of neurons (243-1205 neurons per ∼30 minute-session) and successfully tracked them across days by comparing the distance between neuron ROI centroids (Figure S1A) and verifying that ROIs did not change orientation of their major elliptical axis between sessions (Figure S1B).

**Figure 1:**
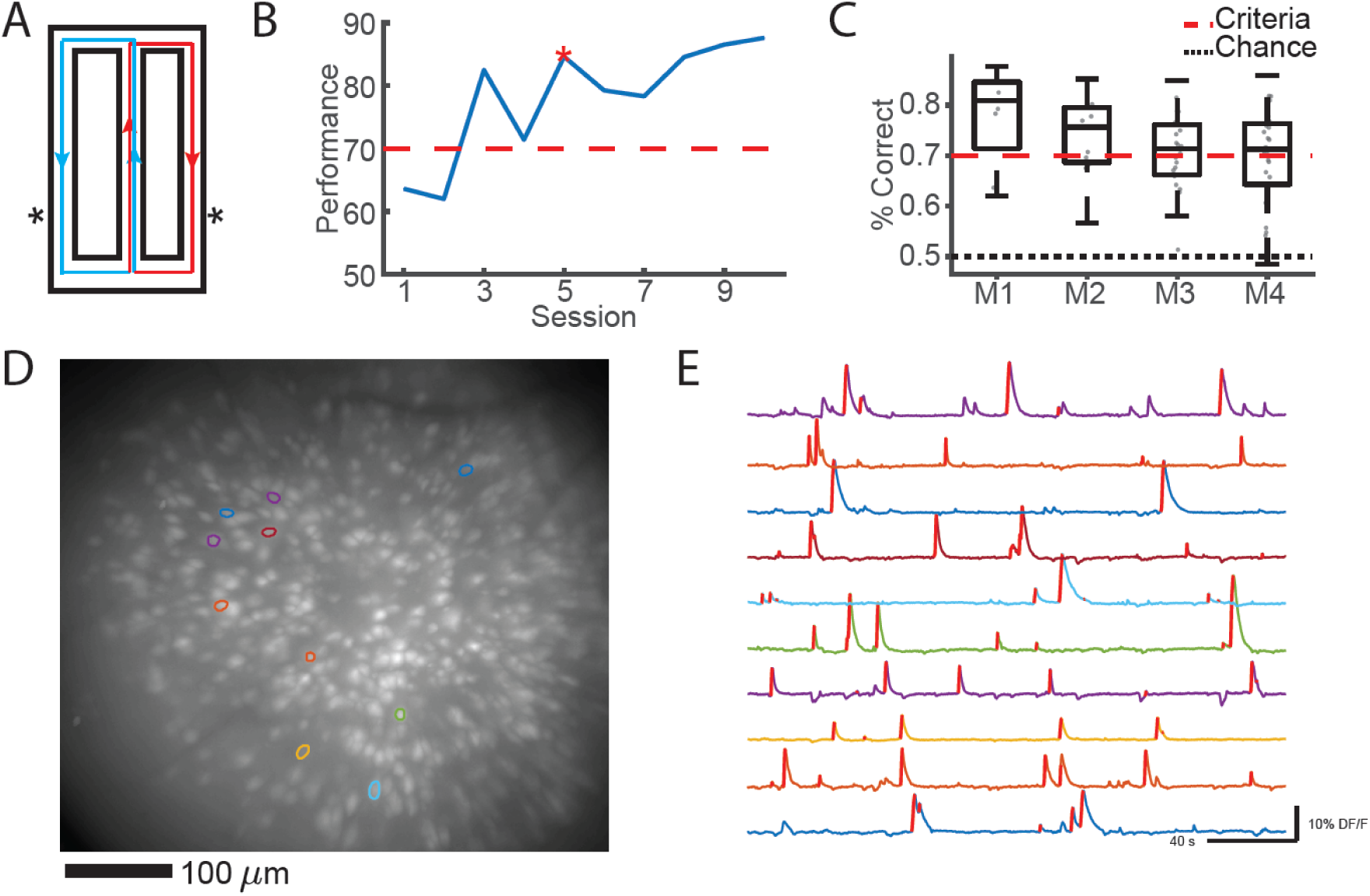
Experimental Setup and Imaging. A) Alternation Maze. Blue = Left turn trajectories, Red = Right turn trajectories, *= location of food reward. B) Example learning curve for one mouse. Red dashed = acquisition criterion (70%), red asterisk = task acquisition day. C) Performance summary for all four mice, all sessions included. Red dashed = criterion, black dashed = chance. D) Example maximum projection from one imaging session with 10 neuron ROIs overlaid. E) Example calcium traces for ROIs depicted in D. Red lines on the ascending phase of each calcium event indicate inferred spiking activity.

### Trajectory-Dependent Activity is Maintained Across Days

The initial studies establishing the existence of trajectory-dependent splitter cells in the hippocampus were performed using electrophysiology in rats (Frank et al., 2000; Wood et al., 2000). Thus, we first wondered if we could detect trajectory-dependent activity in a different species while using a technique with much lower temporal resolution. To do so, we constructed tuning curves representing the probability a given neuron had calcium activity at each spatial bin (1cm) along the stem in correct trials only, and classified neurons as trajectory-dependent splitters if at least 3 bins displayed a significant difference between their tuning curves (p < 0.05, permutation test). We found that we were capable of not only identifying trajectory-dependent cells on a given day (60 ± 23, mean ± s.e.m. across all four mice), but that in many cases these neurons maintained the same functional phenotype across multiple days (Figure 2A-B). Significant trajectory-dependent activity was exhibited by 10% of neurons active on the maze stem across all sessions (12%, 5%, 12%, and 9% for individual mice); note that this method for identifying trajectory-dependent activity is more conservative than that used in previous studies (Frank et al., 2000; Ito, Zhang, Witter, Moser, & Moser, 2015; Wood et al., 2000). Apparent trajectory-dependent activity could also potentially result from factors such as systematic variations in the mouse’s lateral position along the stem. We addressed this in two ways. First, we limited the portion of the maze we considered the stem to exclude any areas where the mouse exhibited stereotypical turning behavior by eye (Figure 2A-B, bottom). Second, we performed an ANOVA for each splitter neuron which included the animal’s upcoming trajectory, position along the stem, speed, and lateral position along the stem as covariates (Wood et al., 2000). We found that a high proportion of our splitter neurons were significantly modulated by upcoming turn direction after accounting for speed and lateral stem position (89%, 72%, 76%, and 83% for individual mice). Together, these results indicate that trajectory-dependent coding exists in mouse CA1 and in many cases maintains the same activity profile across both short and long timescales. To the best of our knowledge, this is the first demonstration of hippocampal trajectory-dependent activity using calcium imaging in mice.

**Figure 2:**
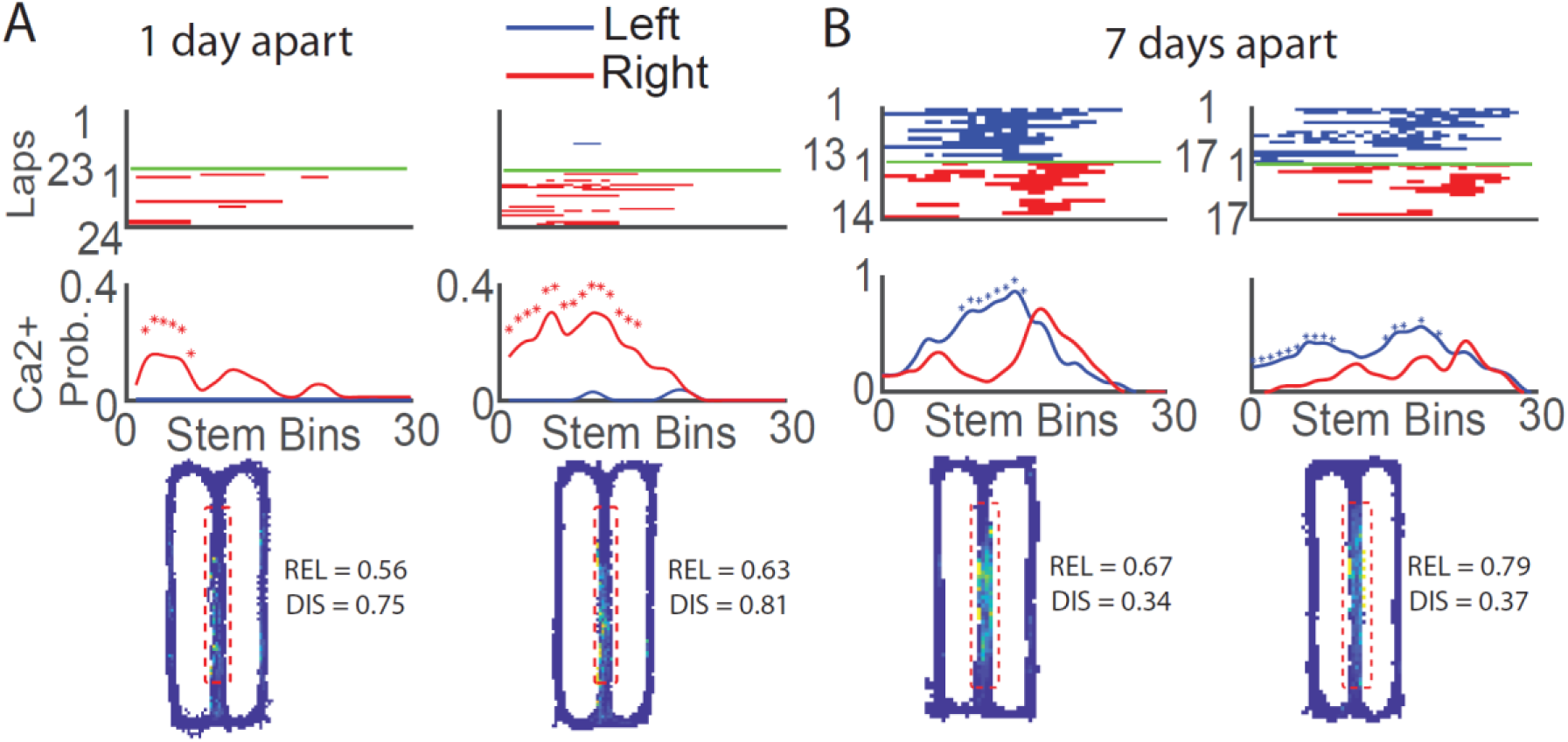
Trajectory-Dependent Activity Persists Across Days. A) Top: Calcium event rasters along the stem for correct trials for two sessions recorded one day apart, sorted by turn direction at the end of the stem. Blue = left, Red = right. Middle: Calcium event probability curves for each turn direction.*p<0.05, shuffle-test. Bottom: occupancy normalized calcium event map with reliability (REL) and discriminability (DIS) scores shown. Red dashed = extent of stem considered in above plots. B) Same as A, but for a different mouse and for sessions 7 days apart.

Additionally, we observed the mean location of spatial firing along the stem progressed backward during the task such that calcium activity occurred at earlier and earlier portions of the stem with time (Figure S2A). This is consistent with a study reporting the backwards-migration of spatial firing with experience (Mehta, Quirk, & Wilson, 2000). Interestingly, we did not find any evidence of consistent migration of spatial firing locations between sessions (Figure S2B).

### Trajectory-Dependent Activity Correlates with Performance

Trajectory-dependent neurons provide information vital to task performance that might be utilized by downstream structures to inform proper motor actions (Albouy et al., 2008; Kahn et al., 2017; Wise & Murray, 1999). This idea is supported by a study which found that trajectory-dependent activity markedly diminished during error trials (Ferbinteanu & Shapiro, 2003). Thus, we predicted that successful task performance would be associated with prominent trajectory-dependent information in the neural code of neurons active on the stem. We utilized two metrics to measure different attributes of trajectory-dependent activity: 1) reliability, which measured the consistency of a cell to fire on its preferred trial type along the entire stem, and 2) discriminability, which measured the magnitude of difference between left and right turn tuning curves along the entire stem (Methods). While most splitter neurons generally had high reliability and discriminability values, neurons with sparser calcium activity for one turn direction could exhibit low reliability and high discriminability (Figure 2A). Conversely, splitter neurons that reliably increased their event rate for one turn direction but still exhibited activity for the other turn direction could have high reliability but low discriminability (Figure 2B).

We obtained significant correlations between each metric and the animal’s performance in a given session (Figure 3A, B top) suggesting that trajectory-information carried in hippocampal neurons might support working memory. To bolster this argument, we also trained a decoder to classify future turn direction using a linear discriminant analysis (LDA) at each spatial bin along the stem based on the neural activity of the population. We found that the accuracy of the LDA decoder positively correlated with the animal’s performance on a given day, which indicated that better separation between upcoming left and right trajectories by the neural code was related to increased memory (Figure 3C, top). These results held for discriminability and LDA accuracy, but not reliability, when we averaged performance across mice (Figure 3A-C, bottom). Last, for each cell, we correlated the left turn and right turn tuning curves and subtracted those values from 1 (1 – ρ) as another metric for trajectory-dependent information. This metric is very conservative because it produces low values (indicating high-trajectory dependent information) for splitter neurons that shift their location along the stem between trial types but not for splitter neurons that modulate event rates in the same location. Despite this we found a significant relationship between performance and 1 – ρ when averaging within mice (Figure 3D, bottom). We obtained similar results when we focused on local metrics of trajectory-dependent activity rather than their average along the entire stem (Figure S3). Together, these results indicate that trajectory-dependent information might facilitate accurate task performance.

**Figure 3:**
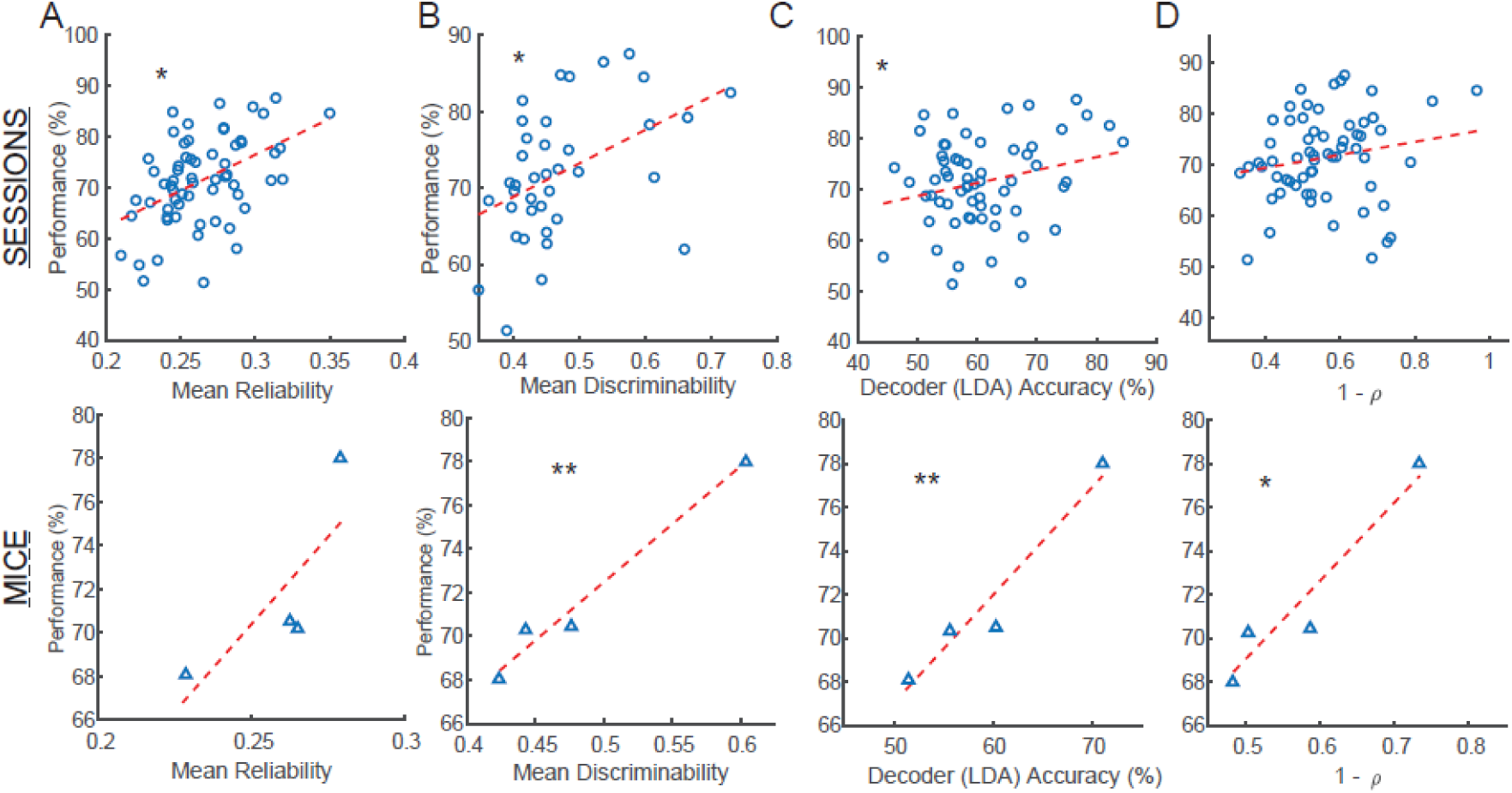
The Quality of Trajectory-Dependent Activity Correlates with Performance. A) Top: Performance for each session versus the mean reliability value for all cells from that session. Circles = all sessions, all mice.*ρ=0.47, p=1.2×10^-4^ Pearson correlation. Bottom: Same as Top but for each mouse, triangles = average for each mouse. B) Same as A, but for the mean discriminability value. *ρ=-0.45, p=0.0043, **ρ=-0.98, p=0.012 (bottom), Pearson correlation. C) Same as A, but for the mean LDA decoder accuracy. *ρ=0.25, p=0.047, **ρ=0.94, p=0.003 Pearson correlation. D) Same as A, but for the mean value of 1 – Spearman’s ρ between left and right tuning curves. *ρ=0.91, p=0.046.

### Trajectory-Dependent Neurons are more likely to Remain Active over Long Timescales than Arm Place Cells

Multiple studies (Cai et al., 2016; Kinsky et al., 2018; Mau et al., 2018; Rubin et al., 2015; Ziv et al., 2013) have shown that hippocampal neurons exhibit significant turnover across days with fewer staying active within the same environment as time progresses. However, these studies all treated the CA1 population as one homogeneous group. Thus, we wondered if splitter neurons, which exhibit highly task relevant information, would be preferentially stabilized within the CA1 network when compared to traditional place cells. As such the hippocampus would maintain a more consistent population of neurons which could be utilized for guiding this behavior. To address this question, we calculated the probability that each pool of neurons remained active in subsequent sessions. We found that splitter cells were more likely to remain active in a later session than arm place cells for both short (Figure 4A) and long (Figure 4B) time lags between sessions. This heightened likelihood that splitter neurons remained active persisted up to 15 days later when considering all mice together (Figure 4C-D) and also held when we compared splitters to place cells with activity on the stem (stem place cells, Figure 4E-F). To mitigate any sampling biases due to the higher event rate of splitter neurons (Figure S4A), we performed an additional analysis where we included only the most active place cells such that their mean event rate matched that of splitters. We obtained similar results when comparing splitter neurons to arm place cells but not stem place cells (Figure S4B-F). Since many stem place cells exhibited trajectory-dependent activity that fell short of meeting our splitter neuron criteria, this suggests that a neuron’s activity level, along with its functional phenotype, also influences whether it stays active on following days. These findings combined support the idea that the strength of task relevant information carried by a neuron influences its likelihood to maintain activity at later time points, which could be exploited for successful memory-guided behavior across days.

**Figure 4:**
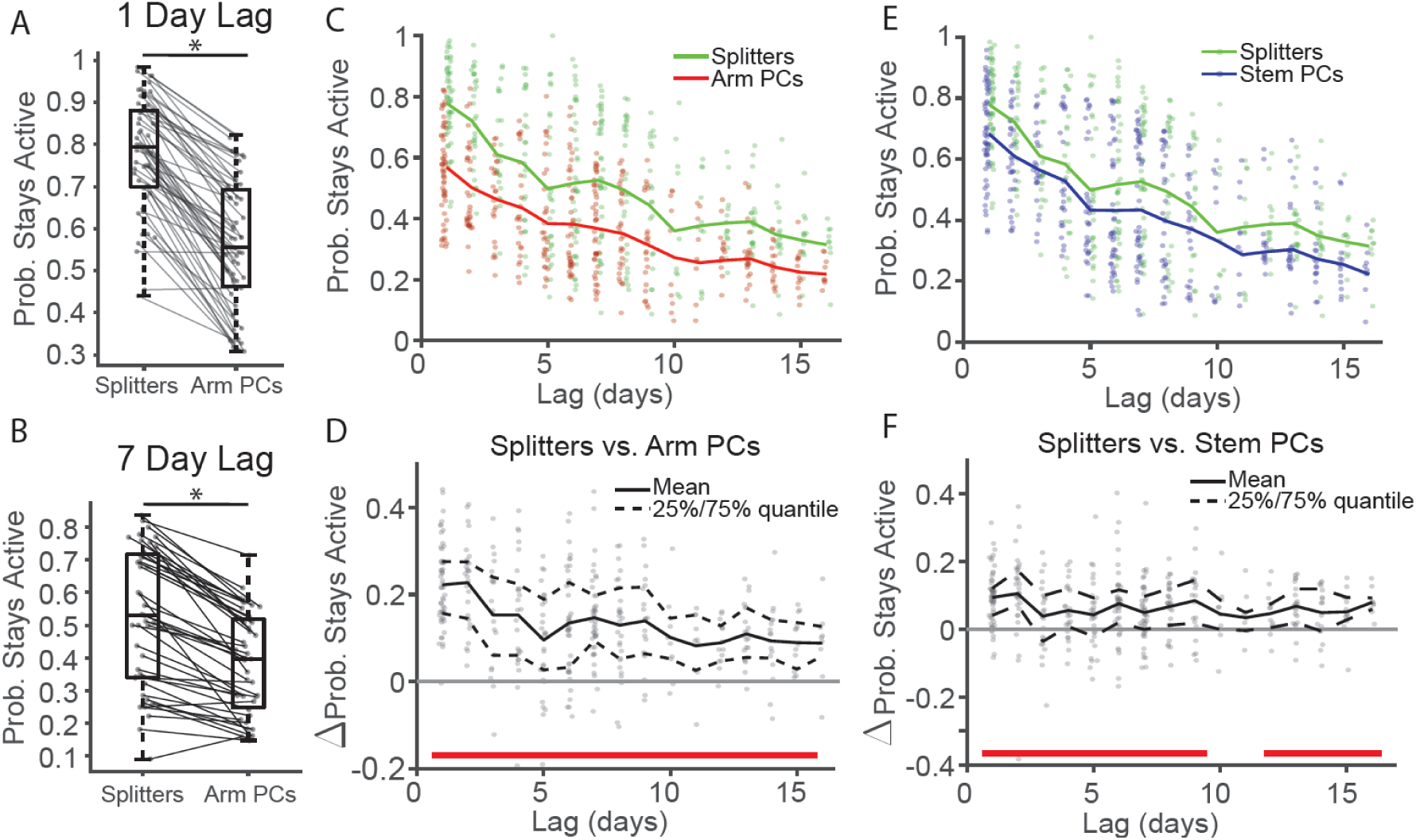
Splitter Neurons are more likely to Remain Active between Sessions than Arm Place Cells. A) Probability splitters and arm place cells (PCs) stay active day later for all mouse. *p=6.4×10^-11^, one-sided signed-rank test. B) Same as B but for all mice and for sessions 7 days apart. *p = 1.1×10^-10^, one-sided signed-rank test. C) Probability splitters and arm place cells stay active versus lag between sessions. Dots: probabilities from individual session-pairs, lines: mean probability at each time lag. Green = splitters, Red = arm PCs. D) Difference between the probability that splitters stay active versus the probability that arm PCs stay active between sessions. Dots: probability differences for individual session-pairs. Black solid/dashed lines: Mean and 25%/75% quantiles of data at each time point. Red bars = significant differences after Holm-Bonferroni correction of one-sided signed-rank test. See Table 1 for signed-rank test p-values at all lags, α = 0.05. E) and F) Same as C-D for splitter neurons vs. stem PCs.

**Table 1:** One-sided Signed-Rank Significance Values for Probability Splitter vs. Place Cells (PCs) Remain Active between Sessions. PCs subsampled to match mean event rate of splitter neurons.

| **Lag (days)** | **1** | **2** | **3** | **4** | **5** | **6** | **7** |
| --- | --- | --- | --- | --- | --- | --- | --- |
| **vs. Arm PCs** | 1.3e-10 | 9.1e-7 | 9.4e-5 | 4.3e-5 | 2.2e-4 | 4.5e-7 | 2.7e-8 |
| **vs. Stem PCs** | 2.e-9 | 5.3e-6 | 0.016 | 1.5e-3 | 0.017 | 5.4e-6 | 1.7e-4 |
| **Lag (days)** | **8** | **9** | **10** | **11** | **12** | **13** | **14** |
| **vs. Arm PCs** | 1.2e-5 | 1.0e-4 | 4.9e-3 | 0.014 | 1.2e-4 | 2.5e-5 | 8.9e-4 |
| **vs. Stem PCs** | 1.1e-4 | 1.7e-3 | 0.082 | 0.097 | 5.2e-3 | 3.3e-4 | 0.012 |
| **Lag (days)** | **15** | **16** |  |  |  |  |  |
| **vs. Arm PCs** | 1.2e-4 | 9.8e-3 |  |  |  |  |  |
| **vs. Stem PCs** | 6.7e-3 | 2.0e-3 |  |  |  |  |  |

We next wondered how the information provided by splitter cells differs from that of other neuron functional phenotypes. To investigate, we decided to compare the long-term spatial coding properties of trajectory-dependent splitter neurons on the stem to return arm place cells (Figure 5A). When examining spatial calcium activity over the entire map across sessions, we found that splitter neurons had a significantly higher 2D event map correlation values than arm place cells (Figure 5B) and that this effect persisted up to 15 days later (Figure 5C) indicating that they were more stable overall. We observed similar results when we compared splitter neurons to stem place cells (Table 2, Figure S5). This indicates that, in addition to staying active over longer time-scales, trajectory-dependent splitter neurons might also better guide memory task performance by providing a more consistent representation of space than place cells.

**Figure 5:**
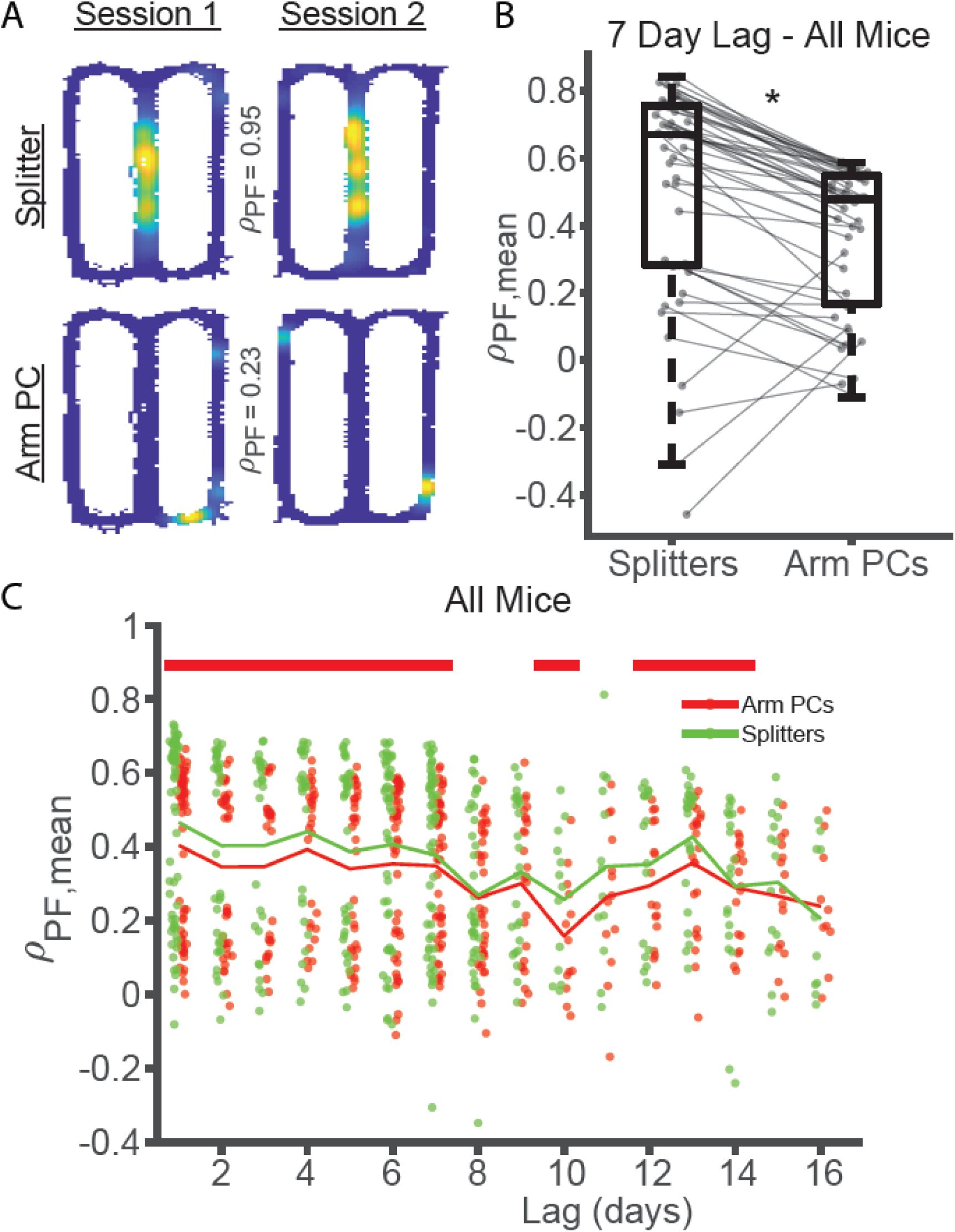
Splitters Maintain More Consistent Spatial Information than Place Cells. A) Example 2D occupancy normalized calcium event maps from the same splitter neuron (top) and arm place cell (bottom) between sessions on adjacent days. The higher Spearman correlation between splitter neuron event maps indicates more consistent spatial activity. B) Mean spatial correlations for splitter neurons versus return arm PCs for all sessions seven days apart from all. *p=4.5×10^-6^, one-sided signed-rank test. C) Mean spatial correlations for splitter neurons and arm PCs versus lag between sessions for all mice/sessions. Red = Arm PCs, green = splitters. Red bars = significant differences after Holm-Bonferroni correction of one-sided signed-rank test, α = 0.05. See Table 2 for raw p-values at all lags.

**Table 2:** One-sided Signed-Rank Significance Values for Mean Spatial Correlation Values of Splitter vs. Arm PCs or Splitters vs. Stem PCs.

| **Lag (days)** | **1** | **2** | **3** | **4** | **5** | **6** | **7** |
| --- | --- | --- | --- | --- | --- | --- | --- |
| **vs. Arm PCs** | 8.6e-8 | 5.0e-4 | 1.7e-3 | 4.6e-3 | 4.5e-3 | 2.6e-4 | 2.6e-3 |
| **vs. Stem PCs** | 4.7e-6 | 4.5e-5 | 7.5e-4 | 5.7e-5 | 5.7e-3 | 3.2e-3 | 1.2e-3 |
| **Lag (days)** | **8** | **9** | **10** | **11** | **12** | **13** | **14** |
| **vs. Arm PCs** | 0.059 | 0.032 | 0.013 | 0.064 | 0.015 | 3.2e-4 | 0.021 |
| **vs. Stem PCs** | 0.23 | 6.2e-3 | 1.2e-4 | 0.024 | 0.59 | 1.4e-3 | 0.44 |
| **Lag (days)** | **15** | **16** |  |  |  |  |  |
| **vs. Arm PCs** | 0.15 | 0.78 |  |  |  |  |  |
| **vs. Stem PCs** | 0.047 | 0.25 |  |  |  |  |  |

### Trajectory-Dependent Neurons Display a Rapid Onset Followed by Stable Activity

Next, we examined the ontogeny of trajectory-dependent behavior. We hypothesized two different scenarios could support the emergence of splitters. In line with a study showing that unstable neurons can support well-learned behavior (Driscoll, Pettit, Minderer, Chettih, & Harvey, 2017), splitters could slowly ramp up/down their splitting behavior or they could come online suddenly and turn off just as suddenly. On the other hand, previous research presented the idea that neurons pre-disposed to become place cells can come online suddenly after a head-scanning/attention event (Monaco, Rao, Roth, & Knierim, 2014), which is potentially supported by the presence of reliable sub-threshold depolarizations of those neurons caused by calcium activity in its dendritic arbor (Bittner, Milstein, Grienberger, Romani, & Magee, 2017; Diamantaki et al., 2018; Lee, Lin, & Lee, 2012; Sheffield & Dombeck, 2014, 2019). In line with this idea, splitter neurons could rapidly develop trajectory-dependent activity and then maintain that activity thereafter. To address this question, we identified the day when each neuron we recorded first exhibited significant trajectory-dependent activity, and then tracked whether that neuron retained trajectory-dependent activity along a similar proportion of the maze stem (splitting extent) in subsequent sessions. We found evidence for heterogeneity in the ontogeny of splitting, with some neurons exhibiting a rapid onset of trajectory-dependent activity (Figure 6A) while others ramped up their trajectory-dependent activity in the days prior to becoming a splitter (Figure 6B). Each onset type appeared to maintain stable trajectory-dependent activity afterward since, for individual mice, splitting extent remained higher in the day following splitter onset when compared to the day preceding splitter onset (Figure 6C). The rapid onset of trajectory-dependent activity and stable maintenance thereafter was readily apparent when examining group data over longer time scales (+/- 10 days, Figure 6D). We obtained similar results for peak reliability and peak discriminability along the stem (Figure 6E-F). In contrast, we observed only a weak trend for reliability and discriminability averaged along the whole stem (Figure S6), supporting the observation that splitter neurons maintained consistent trajectory-dependent activity along a local portion (∼10-20%) of the stem after their onset. We observed a similar trend for place cells using mutual information as a metric of place coding strength (Figure 6G), suggesting that similar rules govern the onset and fate of trajectory-dependent and spatial coding in hippocampal neurons.

**Figure 6:**
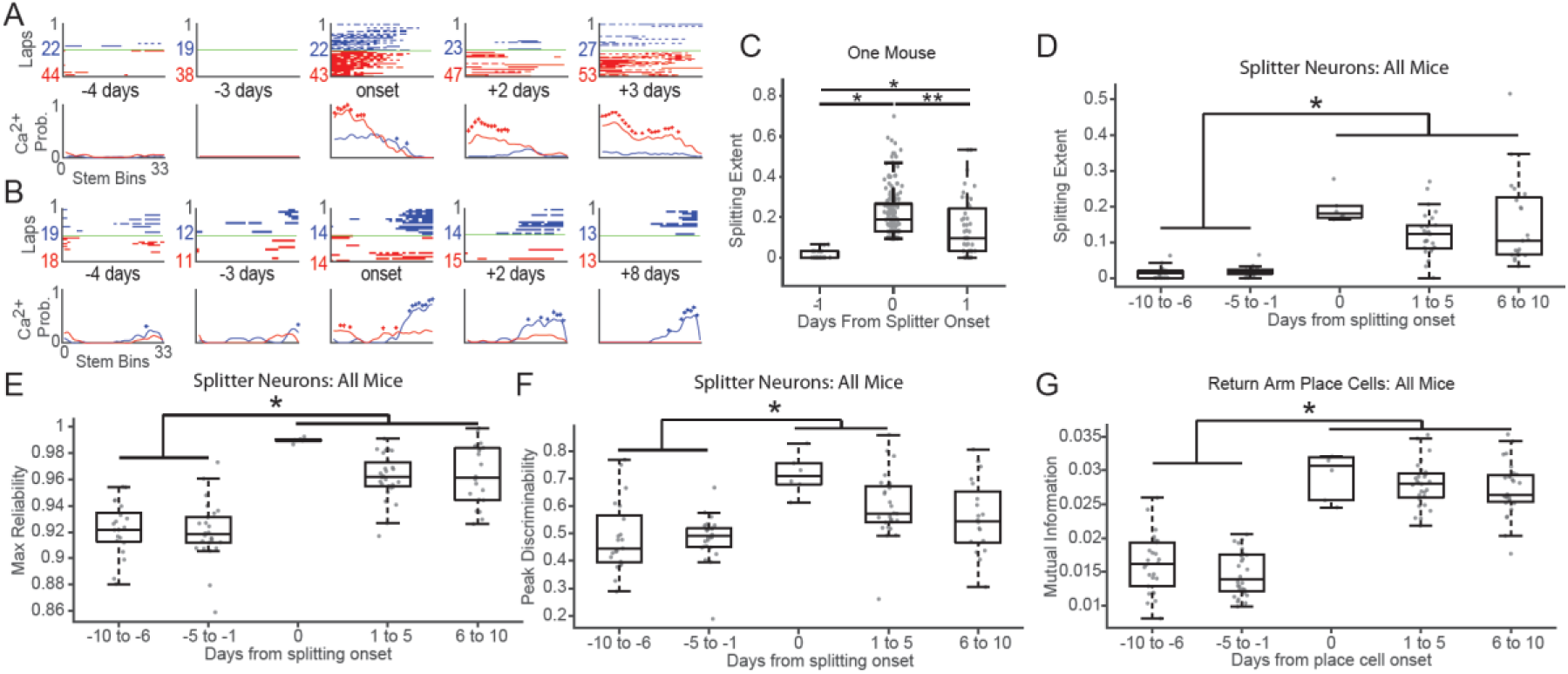
Splitters Come Online Abruptly and Maintain Stable Fields. A) and B) Example splitter across days for two different mice illustrating sudden onset of trajectory-dependent activity followed by stable trajectory-dependent activity thereafter. C) Extent of significant splitting (trajectory-dependent activity) along the stem +/- 1 days from splitter onset for one representative mouse. p = 7.6×10^-19^ Kruskal-Wallis ANOVA, *p < 1.0e-5, **p=0.00025 post-hoc Tukey test. Circles = splitting extent for each neuron. D) Extent of splitting +/- 10 days from splitter onset for all mice. p = 3.1×10^-14^ Kruskal-Wallis ANOVA, *p < 6.0×10^-5^ post-hoc Tukey test. Circles = mean splitting extent of all neurons active on the stem for each session. E) Max reliability score +/- 10 days from splitter onset for all mice. p = 2.8×10^-12^ Kruskal-Wallis ANOVA, *p < 2×10^-4^ post-hoc Tukey test. F) Mean discriminability score +/- 10 days from splitter onset for all mice. p = 4.7×10^-6^ Kruskal-Wallis ANOVA, *p < 0.003 post-hoc Tukey test. G) Mean mutual information +/- 10 days from return place cell onset for all mice. We obtained similar results for stem place cell onset (data not shown). p = 4.7×10^-17^ Kruskal-Wallis ANOVA, *p < 6×10^-4^ post-hoc Tukey test.

### Place Cell Onset Coincides With or Precedes Splitter Onset

We next wondered if hippocampal neurons displayed significant spatial tuning before, during, or after they exhibited trajectory-dependent firing. As shown above, splitter cells produce accurate spatially-modulated activity (Figure 5) and have a similar onset/offset trajectory to place cells (Figure 6); thus, we hypothesized that the onset of trajectory-dependent firing in hippocampal neurons would either coincide with or follow their onset as place cells. To test this idea, we first tallied the onset day of each cell phenotype. We found, while both functional phenotypes were present from day 1 and continued to come online throughout the experiment (Figure 7A,C), the bulk of place cells were recruited on day 1. In contrast, and in agreement with a previous study (Bower, Euston, & McNaughton, 2005), the recruitment of splitter cells did not peak until several days later (Figure 7A,C), suggesting that trajectory-dependent activity tended to emerge more slowly than spatial activity. This could occur independently in two different groups of neurons, or it could occur serially with each neuron first becoming a splitter cell only after becoming a place cell. Thus, to test if this delay in splitter cell ontogeny occurred in the same cells, we directly compared the day a cell became a place cell to the day it began to exhibit trajectory-dependent activity. We found that in the majority of neurons, trajectory-dependent activity onset occurred simultaneously with place field onset, while a different population of neurons exhibited trajectory-dependent activity only after first becoming place cells (Figure 7B,D). Thus, place cells and splitter cells occupy an overlapping population of neurons with spatial responsivity coinciding with or preceding trajectory-dependent coding.

**Figure 7:**
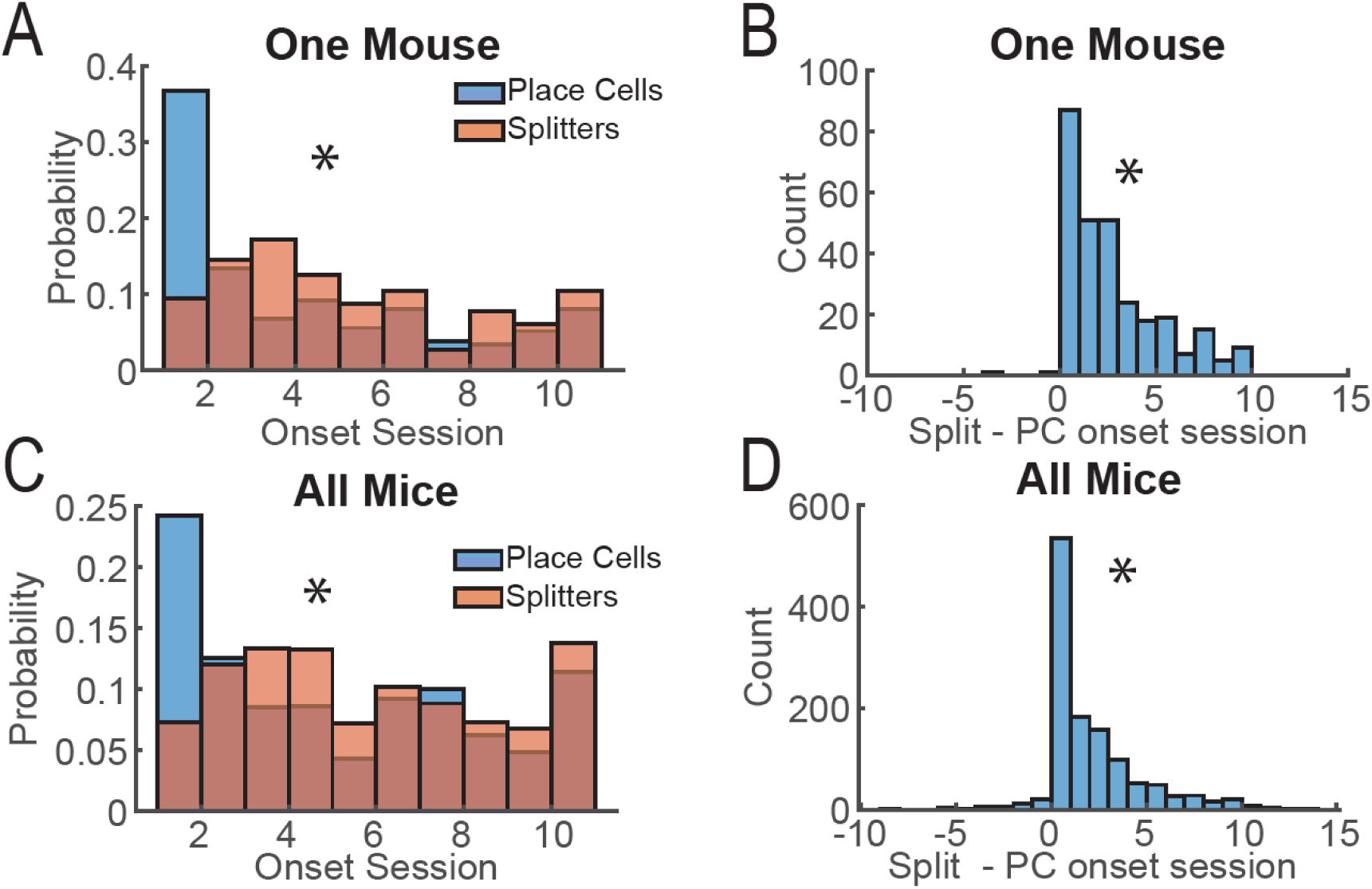
Place Cell Onset Coincides With or Precedes Splitter Onset. A) Histogram of the first (onset) day a neuron exhibits a splitter cell or place cell phenotype for one mouse. *p = 3.7×10^-17^, one-sided Kolmogorov-Smirnov test. B) Difference between splitter cell onset day and place cell onset day for one mouse. *p=2.5×10^-24^ χ2 goodness-of-fit test, mean = 2.3, median = 2. C) Same as A) but for all mice. *p = 2.2×10^-17^, one-sided Kolmogorov-Smirnov test. D) Same as B) but for all mice. *p=5.3×10^-107^ χ2 goodness-of-fit test, mean = 1.6, median = 1.

## DISCUSSION

From an evolutionary perspective, one adaptive function of memory is the ability to provide information vital to survival. Thus, maintaining activity and consistency in neurons encoding information pertinent to survival might provide a mechanism for preferentially strengthening connections with downstream structures via consistent replay of the same sequences (Buzsáki, 2015; Diba & Buzsáki, 2007; Louie & Wilson, 2001; Maboudi et al., 2018; Pfeiffer & Foster, 2013). Conversely, if the pool of neurons available to encode a given memory remains fixed, then forgetting of incidental information through the turnover/silencing of neurons not required for survival is adaptive (Hardt, Nader, & Nadel, 2013) because it could increase the numbers of neurons available to encode other relevant information (Richards & Frankland, 2017). Here, we utilized *in vivo* calcium imaging with miniaturized microscopes to explore this idea (Figure 1) by investigating the development and fate of trajectory-dependent splitter neurons (Frank et al., 2000; Wood et al., 2000) (Figure 2). To the best of our knowledge, this is the first demonstration that trajectory-dependent hippocampal activity exists in mice and that it can be detected with calcium imaging. Since trajectory-dependent splitter neurons contain information relevant to proper task performance (Figure 3, see also Ferbinteanu & Shapiro, 2003), we hypothesized that they would exhibit relatively high stability when compared to other neuron phenotypes.

Several lines of evidence support this hypothesis. First, splitter neurons are more likely to remain active across long time scales than neurons that only provide information about the animal’s current location on the return arm (Figure 4). Second, splitters come online abruptly and then maintain a stable readout of trajectory up to 10 days after becoming a splitter (Figure 6). Splitters also provide a more consistent signal of the animal’s current location than do other neurons (Figure 5), further supporting their long-term stability. Last, we found that splitter cells are a dynamic subpopulation of place cells with the onset of place coding generally preceding the onset of trajectory-dependent activity (Figure 7). This finding concurs with the slow increase of trajectory-dependent activity with experience found in a previous study (Bower et al., 2005). These data combined support the idea that neuron phenotype influences its subsequent stability (Zaremba et al., 2017) and the consistency of the information it provides to downstream structures. More broadly, this study supports the idea that adaptive memories are encoded in a relatively stable subpopulation of neurons, freeing the remaining pool of neurons to undergo plasticity during new learning (Grosmark & Buzsáki, 2016; van de Ven, Trouche, McNamara, Allen, & Dupret, 2016). However, how downstream regions can utilizing a constantly changing landscape of hippocampal inputs to guide behavior remains an open question, as place fields along the stem drift steadily backwards throughout each session (Figure S2) and day-to-day turnover even in relatively stable splitter neurons can still sometimes be quite high (Figure 4C,E).

Our study utilizes single-photon imaging to perform longitudinal tracking of hippocampal neuron activity and confirms existing studies that show increasing turnover of coactive neurons with time (Cai et al., 2016; Rubin et al., 2015; Ziv et al., 2013). However, a recent study performed in songbirds demonstrated that imaging artifacts, specifically small shifts in the z-plane of single-photon imaging, could entirely account for putative cell turnover (Katlowitz, Picardo, & Long, 2018). Thus, the turnover we and others observe in hippocampal neurons could likewise be artefactual. While relevant, this concern is mitigated in our study for a number of reasons. First, the Katlowitz et al. (2018) study was performed in the basal ganglia of songbirds while they performed a stereotyped behavior supported by highly stable firing responses of neurons over short and long timescales (Guitchounts, Markowitz, Liberti, & Gardner, 2013; Hahnloser, Kozhevnikov, & Fee, 2002; Margoliash & Yu, 2009). In contrast, our study was performed in CA1 of the mouse hippocampus, a highly plastic brain region exhibiting complete, monthly turnover of afferent connections (Attardo, Fitzgerald, & Schnitzer, 2015; Pfeiffer et al., 2018) that also exhibits a high degree of drift in neuron firing responses over relatively short time-scales (Mankin et al., 2012; Manns et al., 2007). Second, studies utilizing activity-dependent tagging of neurons also find that the overlap between active cells in the mouse hippocampus declines with time between sessions (Cai et al., 2016; Kitamura et al., 2017), supporting long-term hippocampal cell turnover as a real phenomenon. Most importantly, our study compares the *relative* turnover rates of two different cell phenotypes: splitter cells and place cells. Thus, even if day-to-day misalignments in the z-plane forced neurons out of focus, this would occur equally for both splitters and place cells. Therefore, concerns about imaging artifacts cannot explain our finding that splitter cells are more persistently active across long time scales than place cells.

One notable study found that lesions or optogenetic silencing of nucleus reuniens, an important communication hub between the medial prefrontal cortex and dorsal CA1 of the hippocampus, significantly reduced trajectory-dependent activity in rat CA1 neurons while having no impact on a rat’s performance of a spatial alternation task (Ito et al., 2015). Those results directly challenge our finding that the quality of trajectory-dependent information contained in CA1 activity patterns correlates with a mouse’s performance (Figure 3). One potential reason for this discrepancy is that their intervention only partially reduced trajectory-dependent information without eliminating it, allowing the splitter cells remaining to provide adequate information for proper task performance. In fact, optogenetic silencing of nucleus reuniens produced a smaller deficit in trajectory-dependent activity than did lesions; even lesions eliminated trajectory-dependent activity predicting future trajectories only. Information related to past trajectories, which could be utilized by downstream structures to help make the correct upcoming turn, was maintained. Second, relatively easy tasks might be less resistant to a partial disruption and rats performed at close to ceiling levels in the Ito et al. (2015) study. Our mice performed at lower levels, though still well above chance, indicating that the spatial alternation task might place higher attentional and cognitive demands on mice than on rats. Last, Ito et al. (2015) also utilized the difference in peak firing rate on left versus right trials as a metric for trajectory-dependent activity. This calculation does not account for trajectory-dependent information provided by neurons that maintain similar firing rates, but shift their firing location along the stem between left and right trials (see Figure 2B). Thus, trajectory-dependent neural activity could still be important for proper task performance.

Rodents with hippocampal lesions are capable of performing a continuous alternation task (Ainge, van der Meer, Langston, & Wood, 2007). This raises the question: how important is trajectory-dependent activity if mice can perform the task without the hippocampus at all? We have two responses to this question. First, long-term lesions test necessity, not sufficiency, since these lesions can induce compensatory plasticity that could allow non-hippocampal regions to support the task (Packard & McGaugh, 1996). Second, under normal conditions the hippocampus might still be the default brain region for task performance in spatial alternation. This is emphasized by Goshen et al. (2011), who demonstrated that mice cannot perform long-term recall of a putatively hippocampal-independent contextual fear memory (Bontempi, Laurent-Demir, Destrade, & Jaffard, 1999; Debiec, LeDoux, & Nader, 2002; Frankland, Bontempi, Talton, Kaczmarek, & Silva, 2004; Kim & Fanselow, 1992; Kitamura et al., 2017, 2009; Winocur, Frankland, Sekeres, Fogel, & Moscovitch, 2009) when hippocampal inactivation is limited to a short time period before the task; however, mice became capable of successful long-term memory recall when this inactivation was extended over a long time period prior to performing the task. This study and others (Meira et al., 2018; Sparks, Lehmann, Hernandez, & Sutherland, 2011; Sutherland, O’Brien, & Lehmann, 2008; Wang, Teixeira, Wheeler, & Frankland, 2009; Wiltgen et al., 2010) support the idea that the hippocampus is vital for long-term recall under normal conditions and that redundant pathways are recruited for episodic memory retrieval only if chronic aberrant activity is detected in the hippocampus.

Through what mechanism do trajectory-dependent neurons maintain greater stability across long time-scales? After the initial onset of trajectory-dependent behavior, these neurons could receive feedback from dopaminergic neurons originating in the ventral tegmental area (VTA) during learning (Gomperts, Kloosterman, & Wilson, 2015) or from locus coeruleus (LC) neurons during post-learning sleep (Takeuchi et al., 2016) that could strengthen afferent connections to splitter neurons. This could also occur during sharp-wave ripple related replay of prior trajectories (Diba & Buzsáki, 2007; Pfeiffer & Foster, 2013) in conjunction with simultaneous dopaminergic inputs from VTA (Gomperts et al., 2015). However, this mechanism would also strengthen all cells active en route to the goal location whether they carried information about trajectory or not. One recent study found that trajectories leading to larger rewards were preferentially replayed over trajectories leading to smaller rewards (Michon, Sun, Kim, Ciliberti, & Kloosterman, 2019). Thus, one possibility is that since trajectory-dependent neurons are more useful for predicting how to obtain reward than pure place cells, they might be preferentially reactivated during sharp-wave ripple events, an idea that warrants future testing.

Taken together, our results highlight the influence of cell phenotype on its subsequent stability, and suggest that the emergence of task-related trajectory-dependent coding coincides with or follows the emergence of spatial coding in neurons. Future work should investigate mechanisms supporting the stability and emergence of trajectory-dependent neurons.

## SUPPLEMENTAL FIGURES

**Figure S1:**
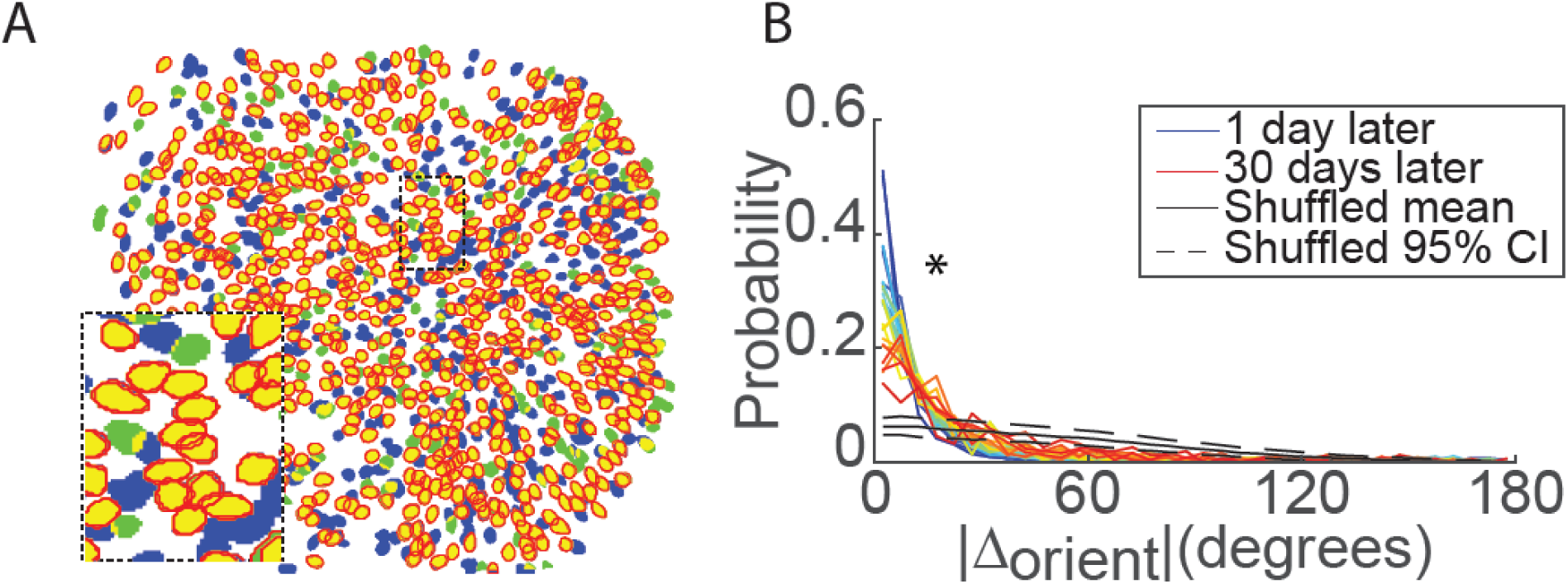
Neuron Extraction and Across-Session Registration. Related to Figure 1. A) Example neuron registration between two sessions. Blue/Green = pixels corresponding to putative ROIs extracted in the 1st/2nd session only. Yellow = pixels corresponding to portions of ROIs active in both sessions. Red = outline of ROIs matched as the same neuron between sessions. B) The small size of changes in ROI orientation between sessions indicate proper neuron registration between sessions.

**Figure S2:**
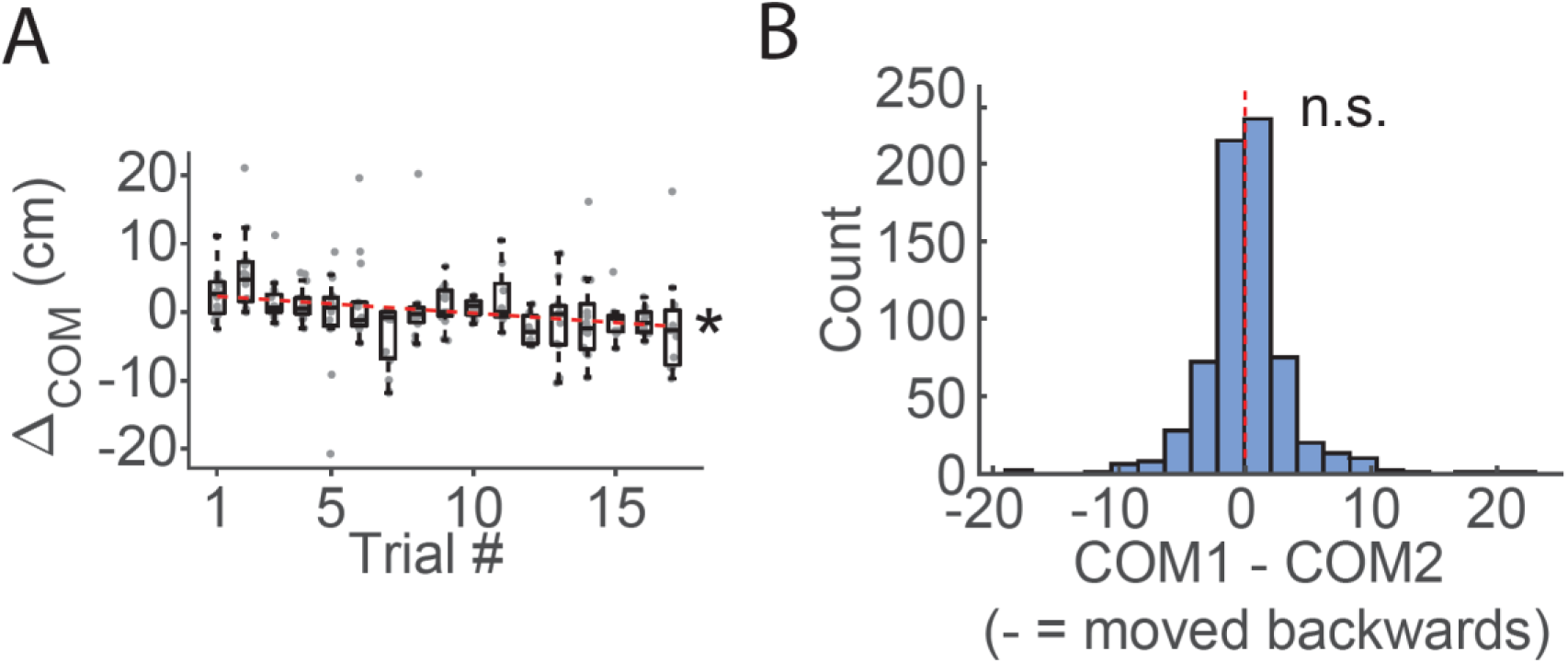
Backward Migration of Spatial Firing Across but not Between Sessions. Relate to Figure 2. A) The centroid of spatial firing on the stem relative to its mean location across the entire session drifts backwards throughout the session. Circles = centroid shifts for each neuron active on the stem. Example session from one mouse for right turns only. *r = −0.28, p=5.1e-5, t = −4.1 for null hypothesis that slope = 0. B) Average change in centroid location between adjacent sessions for all mice between two sessions indicates that place field location does not drift between sessions. p = 0.67 t-test.

**Figure S3:**
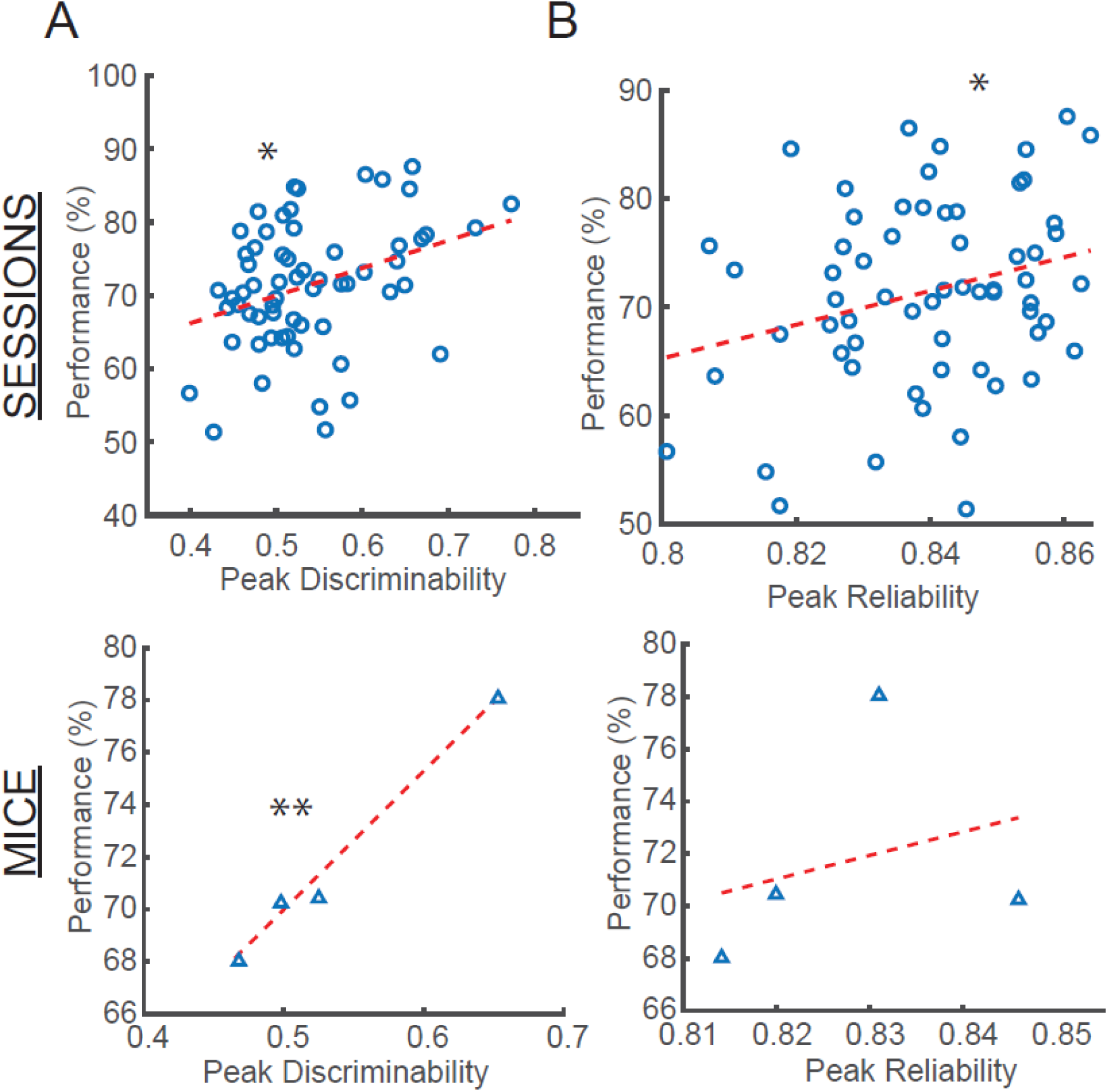
The Quality of Local Trajectory-Dependent Information Correlates with Performance. Related to Figure 3. A) Top: Performance for each session versus the peak discriminability value for all cells from that session. Circles = all sessions, all mice. Bottom: Same as Top but for each mouse, triangles = average for each mouse. *ρ=0.35, p=0.0056, ** ρ=0.99, p=0.0066 Pearson correlation. B) Same as A, but for the peak reliability value. *ρ=-0.27, p=0.031 Pearson correlation.

**Figure S4:**
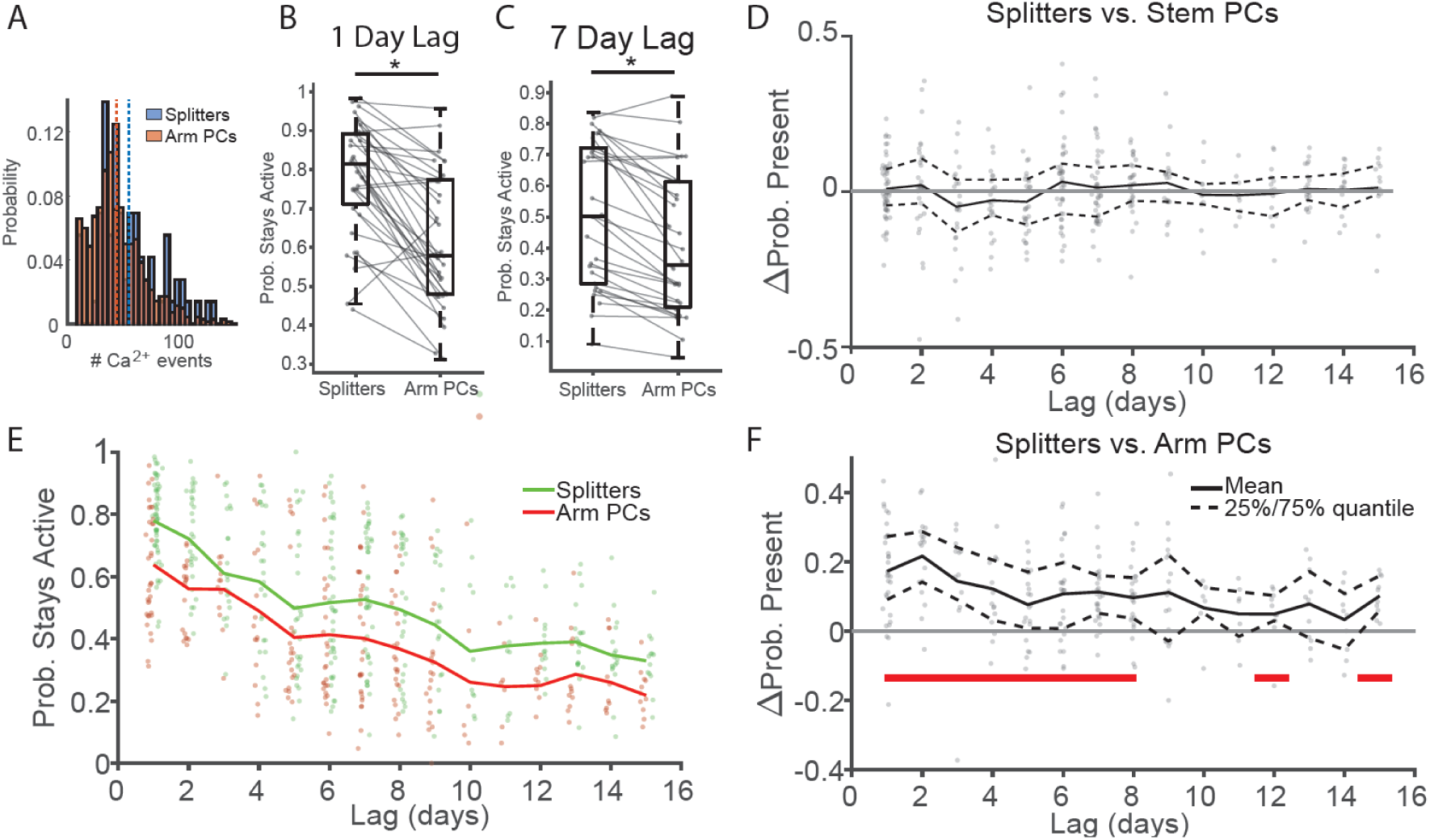
Decreased Turnover Rates for Non Event-Rate Matched Splitters versus Place Cells. Related to Figure 4. A) Histogram from one session showing that Ca2+ event rates are higher for splitters for than for arm PCs. Blue/red dashed: mean number Ca2+ events for splitter/arm PCs. B) Probability splitters and PCs stay active one day later for all mice. *p=3.2×10^-6^, one-sided signed-rank test. C) Probability splitters and PCs stay active seven days later for all mice. *p=3.7×10^-6^, one-sided signed-rank test. D) Difference between the probability that splitters stay active versus the probability stem PC stay active. Dots: probability differences for individual session-pairs. Black solid/dashed lines: Mean and 25%/75% quantiles of data at each time point. See Table 3: One-sided Signed-Rank Significance Values for Probability Splitter vs. Return Arm Place Cells (APCs) are Present, Event-Rate Matched.for one-sided signed-rank test p-values at all lags. E) Probability splitters and arm place cells stay active versus lag between sessions. Dots: probabilities from individual session-pairs, lines: mean probability at each time lag. Green/red: splitters/arm PCs. F) Same as D for splitters vs. arm PCs. Red bars = significant differences after Holm-Bonferroni correction of one-sided sign test, α = 0.05.

**Table 3:**
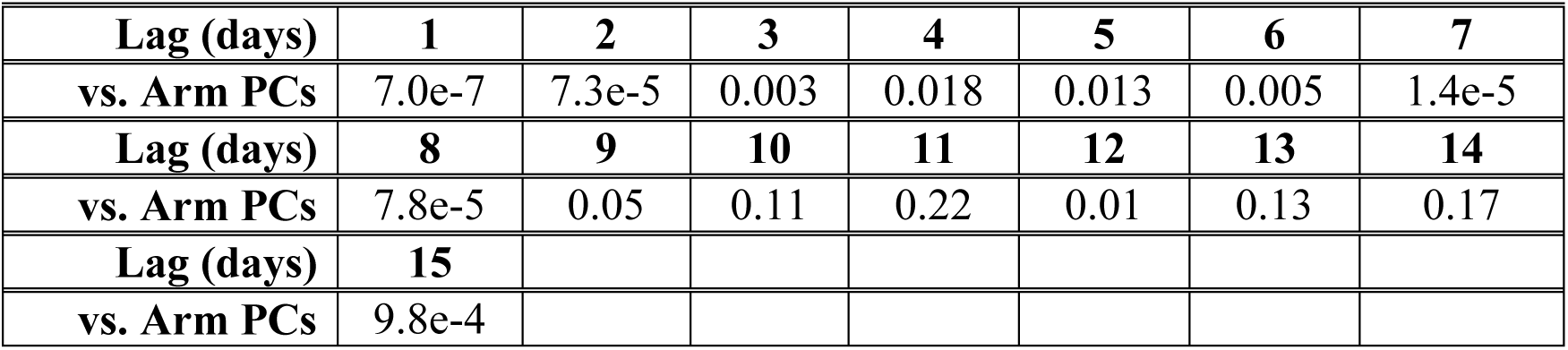
One-sided Signed-Rank Significance Values for Probability Splitter vs. Return Arm Place Cells (APCs) are Present, Event-Rate Matched.

**Figure S5:**
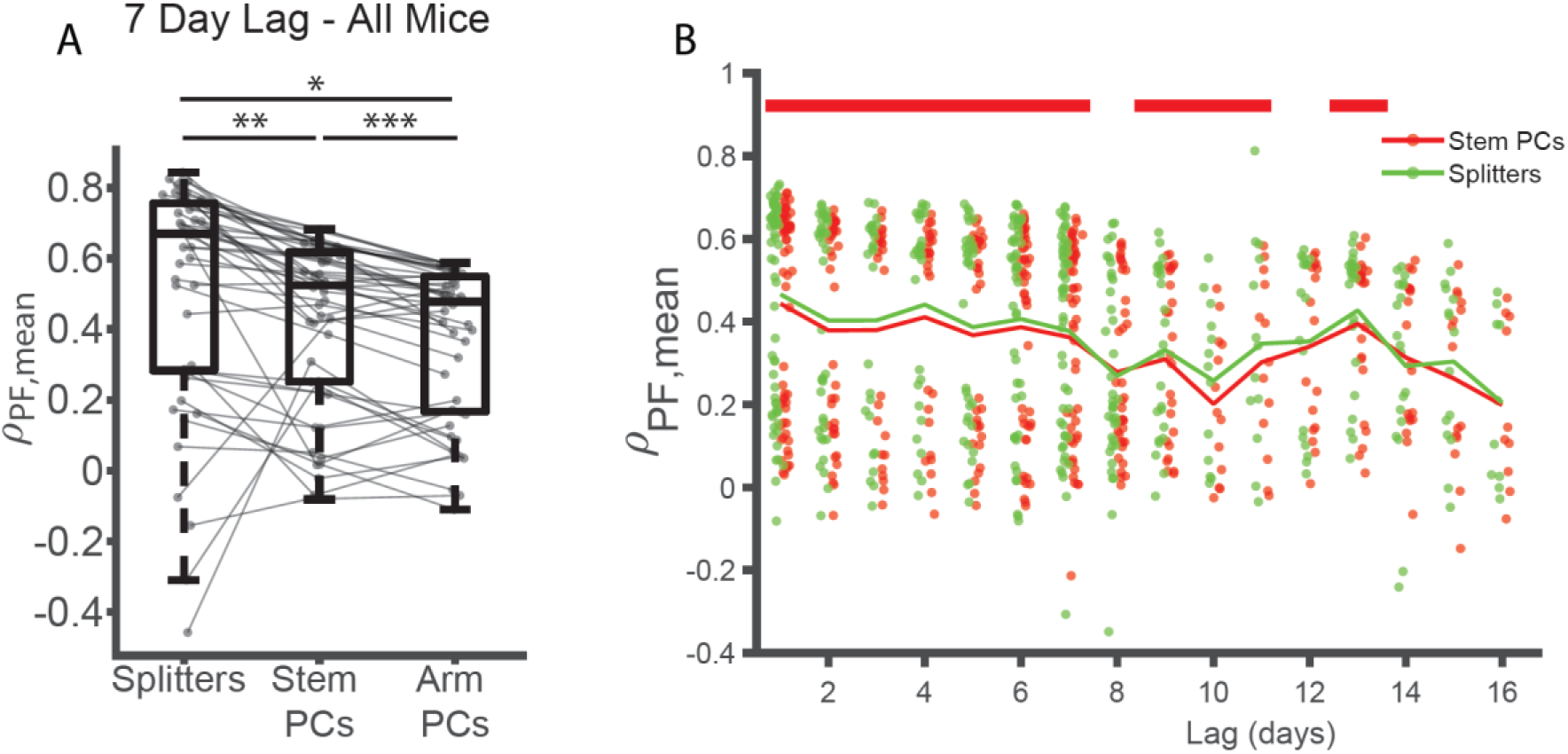
Spatial Consistency of Splitters vs. Stem Place Cells (PCs). Related to Figure 5. A) Mean spatial correlations for splitter neurons versus stem PCs and return arm PCs for all sessions seven days apart from all mice. *p=8.6×10^-6^, **p=4.5×10^-6^, **p=6.5×10^-5^ one-sided signed-rank test. B) Mean spatial correlations for splitter neurons and stem PCs versus lag between sessions for all mice/sessions. Red = Arm PCs, green = splitters, red bars = significant differences after Holm-Bonferroni correction of one-sided sign-test, α = 0.05. See Table 2 for p-values at all lags.

**Figure S6:**
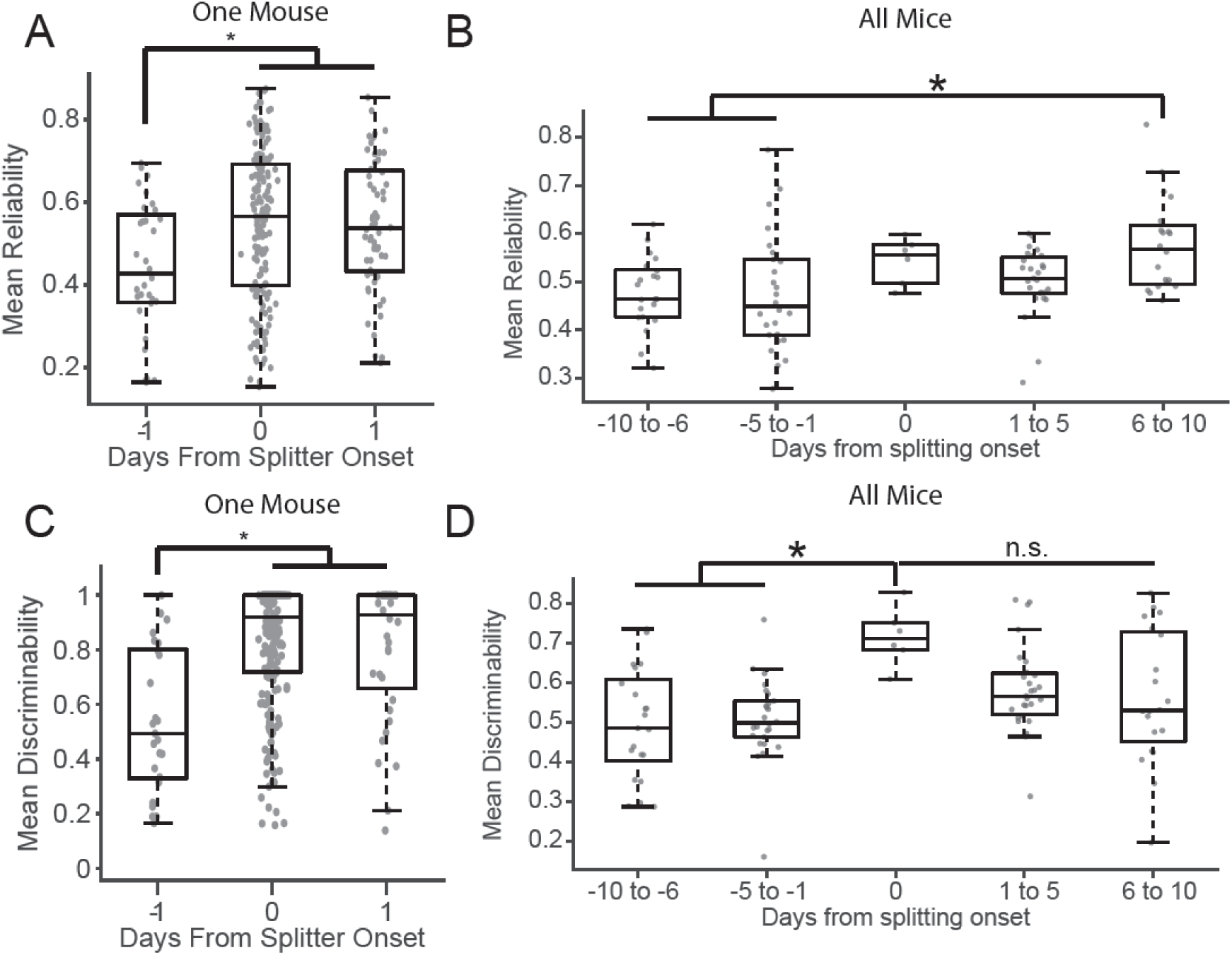
Tracking the Onset of Trajectory-Dependent Activity along the Entire Stem. Related to Figure 6. A) Mean reliability along the stem +/- 1 days from splitter onset for one representative mouse. p = 0.008 Kruskal-Wallis ANOVA, *p < 0.03 post-hoc Tukey test. Circles = mean reliability score for each neuron. B) Mean reliability +/- 10 days from splitter onset for all mice. p = 0.0024 Kruskal-Wallis ANOVA, *p < 0.02 post-hoc Tukey test. Circles = mean of mean reliability of all neurons active on the stem for each session. C) Mean discriminability along the stem +/- 1 days from splitter onset for one representative mouse. p = 1.7×10^-6^ Kruskal-Wallis ANOVA, *p < 0.0004 post-hoc Tukey test. Circles = mean discriminability score for each neuron. D) Mean discriminability +/- 10 days from splitter onset for all mice. p = 0.0014 Kruskal-Wallis ANOVA, *p < 0.006 post-hoc Tukey test. Circles = mean of mean discriminability of all neurons active on the stem for each session.

## METHODS

### Animals

Five male C57/BL6 mice (Jackson Laboratories), age 3-14 months and weighing 25-30g were used. One mouse was excluded from analysis after performing the experiment due to the inability to correct motion artifacts in his imaging videos. Mice were housed socially with 1-3 other mice in a vivarium on a 12hr light/dark cycle with lights on at 7am and given free access to food and water. All mice were singly housed after surgery. All procedures were performed in compliance with the guidelines of the Boston University Animal Care and Use Committee.

### Viral Constructs

We used an AAV9.*Syn*.GCaMP6f.WPRE.SV40 virus from the University of Pennsylvania Vector Core at an initial titer of ∼4×10^13^ GC/mL and diluted it to ∼5-6×10^12^ GC/mL with sterilized 0.05 phosphate buffered saline (KPBS) prior to infusion into CA1.

### Stereotactic Surgeries

All surgeries were performed in accordance with previously published procedures (Kinsky et al., 2018; Resendez et al., 2016) in accordance with the Boston University Animal Care and Use Committee. Briefly, we performed two stereotactic surgeries and one base-plate implant on naïve mice, aged 3-8 months. Surgeries were performed under 1-2% isoflurane mixed with oxygen. Mice were given 0.05mL/kg buprenorphine (Buprenex) for analgesia, 5.0mL/kg of the anti-inflammatory drug Rimadyl (Pfizer), and 400mL/kg of the antibiotic Cefazolin (Pfizer) immediately after induction. They received the same dosage of Buprenex, Cefazolin, and Rimadyl twice daily for three days following surgery and were carefully monitored to ensure they never dropped below 80% of their pre-operative weight during convalescence. In the first surgery, a small craniotomy was performed at AP −2.0, ML +1.5 (right) and 250nL of GCaMP6f virus was injected 1.5mm below the brain surface at 40nL/min. The needle remained in place a minimum of 10 minutes after the infusion finished at which point it was slowly removed, the mouse’s scalp was sutured, and the mouse was removed from anesthesia and allowed to recover.

3-4 weeks after viral infusion, mice received a second surgery to attach a gradient index (GRIN) lens (GRINtech, 1mm x 4mm). After performing an ∼2mm craniotomy around the implant area, we carefully aspirated cortex using a blunted 25ga and 27ga needle under constant irrigation with cold, sterile saline until we visually identified the medial-lateral striations of the corpus callosum. We carefully removed these striations using a blunted 31ga needle while leaving the underlying anterior-posterior striations intact, after which we applied gelfoam to stop any bleeding. We then lowered the GRIN lens until it touched the brain surface and then proceeded to lower it another 50µm to counteract brain swelling during surgery (note that in two mice we first implanted a sleeve cannula with a round glass window on the bottom without depressing an additional 50µm and then cemented in the GRIN lens during base plate attachment). We then applied Kwik-Sil (World Precision Instruments) to provide a seal between skull and GRIN lens and then cemented the GRIN lens in place with Metabond (Parkell), covered it in a layer of Kwik-Cast (World Precision Instruments), and then removed the animal from anesthesia and allowed him to recover after removing any sharp edges remaining from dried Metabond and providing any necessary sutures.

Finally, after ∼2 weeks we performed a procedure in which the mouse was put under anesthesia but no tissue was cut in order to attach a base plate for easy future attachment of the microscope. To do so, we attached the base plate to the camera via a set screw, carefully lowered the camera objective and aligned it to the GRIN lens by eye, and visualized fluorescence via nVistaHD v2.0/v3.0 until we observed clear vasculature and putative cell bodies expressing GCaMP6f (Resendez et al., 2016), then raised the camera up ∼50µm before applying Flow-It ALC Flowable Composite (Pentron) between the underside of the baseplate and the cured Metabond on the mouse’s skull. After light curing we applied opaque Metabond over the Flow-It ALC epoxy to the sides of the baseplate to provide additional strength and to block ambient light infiltration.

### Imagine Acquisition and Processing

Brain imaging data was obtained using nVista HD (Inscopix) v2/v3 at 1440 x 1280 pixels and a 20 Hz sample rate. Two mice were lightly anesthetized (∼60 seconds) to facilitate camera attachment and then given ∼15 minutes to recover prior to any recordings; the camera was attached to the other two mice while they were awake. Prior to neuron/calcium event identification we first pre-processed each movie using Mosaic (Inscopix) software which entailed a) spatially downsampling by a factor of 2 (1.18 μm/pixel), b) performing motion corrections, and c) cropping the motion-corrected movie to eliminate any dead pixels or areas with no calcium activity. We then extracted a minimum projection of the pre-processed movie for later neuron registration. We replaced isolated dropped frames (maximum 2 consecutive frames) with the previous good frame, and in the rare case where more than 2 frames dropped in a row these frames were excluded from all analyses.

### Neuron and Calcium Event Identification

We utilized custom-written, open-source MATLAB software (available at https://github.com/SharpWave/Tenaspis) to identify putative neuron ROIs and their calcium events in accordance with previously published results (Kinsky et al., 2018; Mau et al., 2018). A neuron had to have at least four calcium events in order to be considered active on a given session.

### Across-Session Neuron Registration

We utilized custom-written, open-source MATLAB software (available at https://github.com/nkinsky/ImageCamp) to perform neuron registration across sessions in accordance with previously published results (see Figure S1). We checked the quality of neuron registration between each session-pair in two ways: 1) by plotting the distribution of changes in ROI orientations between session and comparing it to chance, calculated by shuffling neuron identity between session 1000 times, and 2) plotting ROIs of all neurons between two sessions and looking for systematic shifts I neuron ROIs that could lead to false negatives/positives in the registration. During the course of these checks, we noticed the quality of registration between sessions dropped significantly approximately halfway through the experiment for two mice. Thus, we excluded any registrations occurring between the first and second halves of the experiment for these two mice. Furthermore, the second half of the experiment was excluded for these two mice when calculating the absolute onset session of place cells and splitter cells but was included when calculating the relative onset day for each cell type. Several other session-pairs exhibiting poor registrations based on the criteria above were also excluded, though these were rare.

### Behavioral Tracking and Parsing

Behavioral data was recorded via an overhead camera with Cineplex v2/v3 software (Plexon) at a 30Hz sample rate. Cineplex produced automated tracking of the animal’s position by comparing each frame to a baseline image without the animal in the arena. Imaging and behavioral data were synchronized by TTL pulse at the beginning of the recording. Each video was inspected by eye for errors in automated tracking and fixed manually via custom-written MATLAB software. After fixing all erroneous data points, the animal’s position was interpolated to determine his location at each imaging movie time point.

### Histology

Mice were killed and transcardially perfused with 10% KPBS followed by formalin. Brains of perfused mice were then extracted and post-fixed in formalin for 2-4 more days after which they were placed in a 30% sucrose solution in KPBS for 1-2 additional days. The brains were then frozen and sliced on a cryostat (Leica CM 3050S) in 40 μm sections after which they were mounted and coverslipped with Vectashield Hardset mounting medium with DAPI (Vector Laboratories). We then imaged slides at 4x, 10x, and 20x on a Nikon Eclipse Ni-E epifluorescence microscope to verify proper placement of the GRIN lens above the CA1 pyramidal cell layer.

### Experimental Outline

After recovery from surgery, mice were food deprived to maintain no less than 85% of their pre-surgery weight. Mice were subsequently exposed to a variety of arenas in order to habituate them to navigating with the camera attached. Prior to training on the alternation task, all mice were given 1-4 habituation sessions on the alternation maze. The maze floor (inner dimension = 64 x 29 cm) and walls (height = 18cm) were constructed from 3/8 inch (0.95cm) thick plywood and the barriers between arms were constructed from two 53cm long 1.5 x 5.5 inch (3.8 x 14 cm) pine framing studs. The finished maze consisted of a central stem and two return arms, each 7.5cm wide with 5.7cm wide openings at each of the central stem through which mice could exit or enter the return arms. Two food wells ∼ 0.25 cm deep were created toward the end of each return arm to hold chocolate sprinkles: they were centered 12.5 cm from the end of the maze where mice exited the return arm/entered the center stem. Food was placed in these wells through a small opening in the side of the maze. The arena was sealed with urethane prior to exploration.

Three of the mice were first trained to loop on each side of the maze independently in 3-10 minutes blocks by blocking off access to the other side with Plexiglas dividers in order to familiarize mice with the general task demands, arena, and location of food reward (chocolate sprinkles); the other mouse received one habituation session where he was allowed to freely traverse the maze. Following habituation, mice were placed in the center stem and rewarded at the well on the reward arm regardless of the first turn direction. On subsequent trials, mice were only rewarded if they turned the opposite direction of the previous trial. Mice were allowed to run freely and were only blocked when they a) attempted to reverse course on the central stem, b) attempted to exit the return arm after they had committed to it, or c) attempted to run straight across to the other arm without turning down the central stem after obtaining reward. A mouse was considered committed to an arm after his tail entirely crossed from the edge of the central arm into the stem. Mice generally ran ballistically down the center stem and were allowed to pause once they entered the return arm and after they obtained reward. Food reward was only delivered once the mouse had committed to a return arm in order to avoid providing an auditory cue of reward location. Two mice were forced to alternate in a subset of sessions/trials. One mouse encountered a lapse in performance mid-way through the experiment and began perseverating on one turn direction in blocks: he was subsequently given a number of trials at the beginning of each session where he was forced to turn each direction by blocking off one turn direction with a Plexiglas divider, after which he was then allowed to freely choose turn directions. The other mouse was initially forced to alternate at the end of his habituation looping sessions. All forced trials were not considered during later data analysis. Mice received 1-2 sessions per day. Sessions were terminated each day after 30 minutes or when the mouse stopped consistently running ballistically down the center arm, whichever came first. The experiment lasted 27, 16, 29, and 36 days for the four mice involved.

### Place Cell Identification

Place cell identification was performed as described in Kinsky et al.(2018).

### Trajectory-Dependent/Splitter Cell Identification

Prior to performing any analysis, each mouse’s trajectory data was aligned to that from the first habituation session. This was done by 1) manually rotating the data to correct for any day-to-day changes in maze angle relative to the recording camera, 2) calculating the edges of the mouse’s trajectory as the data points located at the 2.5% and 97.5% points in the cumulative density function of his x/y position data, and 3) adjusting the data by applying the necessary translation and scaling (minimal) to overlay each session’s trajectory on the first session. After aligning data across sessions, the mouse’s trajectory on each trajectory was parsed into his progression through the different sections of the maze, starting at the a) base, then moving down the b) center stem into the c) choice point, then turning into the d) left/right entry to the e) return arm, and finally entered the f) approach to the center stem just after the reward port. The center stem portion was manually identified for each mouse as the point where the mouse’s trajectory into/out of each return arm stopped diverging. This was done in order to mitigate the possibility that trajectory-depending activity was controlled entirely by stereotyped sensory inputs, e.g. the mouse hugging/whisking the left side of the center stem after right turn trial.

After parsing the animal’s behavior into these sections, the center stem was broken up into ∼1cm bins and the event rate for each neuron was calculated for each trial. Tuning curves for each trial type (left or right turn) were then constructed, which consisted of each neuron’s mean event rate for all correct trials at each spatial bin. The difference between these curves was then calculated. To assess significance, we again constructed tuning curves for left/right trials and calculated their difference, but after randomly shuffling trial turn identity 1000 times to establish the likelihood the observed difference between tuning curves could emerge by chance. We then defined splitters/trajectory-depending cells as neurons which had at least three bins whose real tuning curve difference exceeded 950 of shuffled values. In order to exclude spurious identification of splitters we only included neurons that produced a calcium event on the stem of the maze on at least 5 trials.

We calculated several different metrics to quantify the level of trajectory-dependent activity in each neuron. First, we calculated discriminability by summing the absolute value of the difference between tuning curves along all stem bins and then dividing by the sum of tuning curves along all stem bins. Second, we calculated reliability in the following manner: a) we shuffled trial identity 1000 times and calculated the difference between shuffled tuning curves, then b) calculated the proportion of shuffles in which the real difference between tuning curves exceeded that of shuffled, then c) calculated reliability as the mean of this proportion along all the stem bins. Note that splitter neurons by definition must have at least three bins with a reliability value above 0.95 (see above). Last, we calculated the correlation between left and right tuning curves. Note that this metric is very conservative since it produces low correlations for splitters who shift the location of their peak activity between left and right trials along the length of the stem (Figure 2B) but not for splitters who modulate their event rate in the same place along the stem (Figure 2A).

In order to check the robustness of our results and control for any trajectory-dependent information resulting from stereotyped deviations in speed or lateral position along the stem, we also performed an additional analysis in line with previous studies (Ito et al., 2015; Wood et al., 2000). To do so, we first divided the stem into five bins and calculated the average transient probability in each bin for all trials. We then performed an ANOVA analysis using the anovan function in MATLAB for each trajectory-dependent splitter neuron we detected. We used trial type (left/right), stem bin, stem bin x trial type, speed, and the animal’s lateral position as covariates and mean transient probability as our dependent variable. Finally, we considered any neuron to be a trajectory-dependent splitter neuron if had had a significant effect of trial type or trial type x stem bin after accounting for speed and lateral position.

### Linear Discriminant Decoding Analysis

A linear discriminant decoder was trained on data from 50% of trials on a given session using the fitdiscr function in MATLAB. Calcium event activity for each neuron at each time point when the mouse was on the center stem were used as the input variables and the mouse’s upcoming turn direction was used as the response variable. Only correct trials were considered for training. The decoder was then used to predict the turn direction of the other 50% of trials, after which the process was repeated 999 times using a different random 50% of trials for training/decoding. The decoding accuracy was then calculated in ∼3.3cm bins along the stem, and the mean accuracy across all bins was taken as the decoding accuracy for that session.

### Functional Phenotype Designation and Analysis of Neurons that Remain Active between Sessions

We first performed neuron registration between sessions and classified neurons as staying active if they were identified by our cell extraction algorithm on both sessions and produced at least five calcium events (while the mouse was running) through the course of the first recording session being considered in the registration. We then categorized cells into three different functional phenotypes, 1) context-dependent splitter cells, 2) arm place cells, and 3) stem place cells. Splitter cells were designated based on the criteria listed above. Neurons that produced no calcium activity on the stem of the maze and met our place cell criteria were defined as return arm place cells. Neurons that produced calcium activity on the stem and met our place cell criteria but not our splitter neuron criteria were designated as stem place cells. In order to ensure sufficient precision in calculating the probability a functional phenotype stayed active, a session-pair was excluded from analysis if there were fewer than ten cells in either category in the first session being registered. This analysis was performed in two ways: 1) including all cells found for each phenotype, and 2) matching mean event rate between neuron phenotype by excluding the lowest firing rate place cells. In the rare event that place cells had a higher mean firing rate than splitter cells, no place cells were excluded.

### Phenotype Ontogeny Analysis

We tracked the ontogeny splitter cells in three steps. First, we registered all the neurons we recorded across the entire experiment. Second, we identified the first day/session that a neuron passed our statistical criteria to be considered a splitter and defined that session as its onset. Finally, we calculated multiple metrics for the quantity of trajectory-dependent activity produced by each of these neurons (see 0 above) in all the sessions preceding and following onset, excluding any sessions that occurred on the same day. The methodology for tracking place cell onset was identical, except mutual information was used as a metric of spatial information provided by each cell.

## AUTHOR CONTRIBUTIONS

Conceptualization: NRK. Methodology: NRK. Software: NRK, WM, DJS, SJL. Validation: NRK. Formal Analysis: NRK, WM. Investigation: NRK, WM. Resources: NRK, MEH. Data Curation: NRK, WM, EAR. Writing – original draft preparation: NRK. Writing – review and editing: NRK, WM, DJS, EAR, SJL, MEH. Visualization: NRK, WM. Supervision: MEH, NRK. Project administration: NRK, MEH. Funding Acquisition: MEH.

### ACKNOWLEDGEMENTS

First and foremost we would like to thank Dr. Howard Eichenbaum for his help in all stages of this project. We would like to also thank Dr. Jon Rueckemann and Dr. Nick Robinson for input on data analysis, interpretation, and feedback during manuscript preparation. We would like to thank Dr. Steve Gomperts, Dr. Annabelle Singer, Dr. Amar Sahay, Dr. Kamran Diba and the members of their labs for helpful comments during discussions of this work. We would also like to thank Dr. Ian Davison, Dr. Steve Ramirez, Dr. Ehren Newman, and Dr. Mark Kramer, as well as Dr. Andrew Alexander, Dr. Jake Hinman, Dr. John Bladon, Dr. Ryan Place, Dan Sheehan, Winny Ning, Dan Orlin, Jiawen Chen, and Catherine Mikkelsen for feedback and critiques during early stages of data analysis and writing. We would like to thank Annalyse Kohley, Hanish Polavarapu, and Scott Bovino for help with animal training and behavioral tracking quality control. We would like to thank Denise Parisi, Dr. Shelley Russek, and Sandra Grasso for administrative support. We would like to acknowledge the GENIE Program, specifically Vivek Jayaraman, Ph.D., Douglas S. Kim, Ph.D., Loren L. Looger, Ph.D., Karel Svoboda, Ph.D. from the GENIE Project, Janelia Research Campus, Howard Hughes Medical Institute, for providing the GCaMP6f virus. Finally, we would like to acknowledge Inscopix, Inc. for making single-photon calcium imaging miniscopes widely available, and specifically Lara Cardy and Vardhan Dani for all their technical support throughout and after the experiment.

